# Enzyme-constrained models predict the dynamics of *Saccharomyces cerevisiae* growth in continuous, batch and fed-batch bioreactors

**DOI:** 10.1101/2021.07.22.453334

**Authors:** Sara Moreno-Paz, Joep Schmitz, Vitor A.P. Martins dos Santos, Maria Suarez-Diez

## Abstract

Genome-scale, constraint-based models (GEM) and their derivatives are commonly used to model and gain insights into microbial metabolism. Often, however, their accuracy and predictive power are limited and enable only approximate designs. To improve their usefulness for strain and bio-process design, we studied here their capacity to accurately predict metabolic changes in response to operational conditions in a bioreactor, as well as intracellular, active reactions. We used flux balance analysis (FBA) and dynamic FBA (dFBA) to predict growth dynamics of the model organism *Saccharomyces cerevisiae* under different industrially relevant conditions. We compared simulations with the latest developed GEM for this organism (Yeast8) and its enzyme-constrained version (ecYeast8) herein described with experimental data and found that ecYeast8 outperforms Yeast8 in all the simulations. EcYeast8 was able to predict well-known traits of yeast metabolism including the onset of the Crabtree effect, the order of substrate consumption during mixed carbon cultivation and production of a target metabolite. We showed how the combination of ecGEM and dFBA links reactor operation and genetic modifications to flux predictions, enabling the prediction of yields and productivities of different strains and (dynamic) production processes. Additionally, we present flux sampling as a tool to analyze flux predictions of ecGEM, of major importance for strain design applications. We showed that constraining protein availability substantially improves accuracy of the description of the metabolic state of the cell under dynamic conditions. This therefore enables more realistic and faithful designs of industrially relevant cell-based processes and, thus, the usefulness of such models

## 1. Introduction

One of the goals of biotechnology is the design of cell factories to produce metabolites of industrial interest. Metabolic engineering introduces heterologous pathways and rewires cell metabolism aiming to increase product yield, titer and productivity [1]. However, although the production capacity of microorganisms is affected by many external factors such as temperature, pH, oxygen and carbon availability, these interactions are often underestimated during the strain design process. The lack of a strong link between initial strain design and industrial deployment causes the so called “Valley of Death”, where only one in 5,000 to 10,000 innovations make the long route from initial finding to market implementation [2]–[4]. Models of microbial metabolism are increasingly being used to aid the design and steering of bio-processes in an attempt to navigate the “Valley of Death”. We studied the capacity of these models to provide accurate predictions of intracellular active fluxes, key to guide metabolic engineering strategies. Besides, we tested their ability to link strain and bio-process design analyzing how modifications in the reactor environment impact predictions on cell metabolism.

Genome-scale metabolic models (GEM) are mathematical representations of cell metabolism able to establish genotype-phenotype relationships linking genes and enzymes with metabolic reactions. These models are based on annotated genomes and thermodynamics, and information regarding enzyme kinetics and genetic regulation is not required [5]–[7].

The incorporation of resource allocation strategies such as maximum membrane surface area or cell volume to GEM has improved flux predictions [8]. Sánchez et al. introduced the GEM with Enzymatic Constraints using Kinetic and Omics (GECKO) framework to generate enzyme constrained models (ecGEM) by adding additional constraints linked to the limited enzyme production capacity of the cell. In these models, protein abundance and enzyme turnover values (*k_cat_*) limit the flux of the corresponding reactions [9]. The ecGEM of *Saccharomyces cerevisiae* was shown to allow a more extensive and accurate simulation of microbial physiology including overflow metabolism, stress responses and consumption rates of different carbon sources.

Flux balance analysis (FBA) is the most common method to simulate genome-scale metabolism. It uses linear programming to optimize an objective function and has extensively been used to predict cellular growth, product secretion patterns and to develop overproduction strains [6], [7], [10], [11]. FBA assumes time-invariant extracellular conditions consistent with chemostat operation. Still, industrial-scale production is often achieved with batch and/or fed-batch cultures were extracellular conditions vary in time. Therefore, dynamic FBA (dFBA) extends FBA by introducing kinetic equations for extracellular metabolites and biomass. dFBA has been applied to simulate *Escherichia coli* industrial fermentations, compare ethanol production of different *S. cerevisiae* strains during fed-batch growth and identify industrially relevant bottlenecks for ethanol production from xylose[12]–[14]. Whereas FBA only captures one of the multiple solutions that leads to the optimization of the desired objective, sampling algorithms provide distributions of feasible flux solutions that represent the whole feasible flux space. Besides, the establishment of an objective function, which may introduce bias on the predictions, is not required [15].

We used FBA and dFBA to predict growth dynamics of *S. cerevisiae* under industrially relevant conditions and compared simulations using the GEM (Yeast8) and the ecGEM (ecYeast8) with experimental data. We tested the capacity of the models to predict changes in cell metabolism (substrate uptake, growth and product secretions) in response to the operation of the reactor, constituting one of the few examples of combination of ecGEM and dFBA. For the first time, we used flux sampling of ecYeast8 to evaluate central carbon metabolic fluxes at a range of growth rates representative of chemostat, fed-batch and batch growth of *S. cerevisiae*. We tested whether constraining the enzyme production capacity of the cell is enough to limit the flux solution space and improve predictions of intracellular active reactions, of major importance for strain design applications. We provide a set of scripts to easily implement dFBA on traditional and ecGEM as well as a validation dataset containing fermentation-related data of *S. cerevisiae* cells growing in chemostat, batch and fed-batch reactors.

## 2. Materials and methods

Yeast8 and ecYeast8 models were downloaded from https://github.com/SysBioChalmers/yeast-GEM [16]. Model simulations were performed using Python 3.6, COBRApy (version 0.18.1) and glpk as solver [17]. Experimental data used in this study, functions developed for chemostat, batch and fed-batch simulations as well as an example on their use are available at https://gitlab.com/saramorenopaz/ecmodels-predict-growth-dynamics-s.-cerevisiae.git.

### 2.1. EcYeast8 model modifications

#### 2.1.1. Model re-scaling

In ecModels enzymes are treated as metabolites which stoichiometric coefficient is 1/*k_cat_*. In ecYeast8 *k_cat_* values expand 10 orders of magnitude (1 to 10^10^ *h*^-1^) resulting in stoichiometric coefficients below solver tolerance (10^-6^) and numerical instability of the model during flux sampling. To minimize the problem the range of *k_cat_* values was reduced so the maximum allowed *k_cat_* value was 10^6^. Besides, all 1/*k_cat_* coefficients and the protein pool exchange upper bound were scaled by 10^3^ to reduce the impact of rounding errors on flux predictions through enzyme usage reactions. These modifications did not change flux predictions as the contribution of extremely efficient enzymes (i.e. k_cat_ > 10^6^) to the protein pool is negligible.

#### 2.1.2. Model constraints

For all the simulations the upper bounds of exchange reactions to produce acetaldehyde, 2,3-butanediol, glycine, acetate and pyruvate were constrained to match experimental measurements [9]. Also, the transport of Ser from mitochondria to cytoplasm (r_2045_REV) and the cytoplasmic NADP^+^ dependent conversion of isocitrate to 2-oxoglutarate (r_0659No1) were blocked as described in Sánchez et al. [9]. Besides, the reversible transaldolase reaction (r_1048 REVNo1), the reversible reaction of malate dehydrogenase in the cytoplasm (r_0713_REVNo1), the fumarate reductase reaction in the cytoplasm (r_1000No1), the reversible isocitrate dehydrogenase reaction in the cytoplasm (r_0659_REVNo1) and the glutamate decarboxylase reaction (r_0469No1) were blocked by constraining to zero their upper and lower bounds.

### 2.2. Model simulations

#### 2.2.1. Glucose limited chemostat simulations

During chemostat simulations metabolic fluxes were calculated setting the bounds of the biomass reaction (r_2111) equal to the dilution rate (D, *h*^-1^) and minimizing glucose consumption as objective for FBA optimization (maximize r_1714 for Yeast8 and minimize r_1714_REV for ecYeast8) [18]. The dilution rate was varied from 0.05 *h*^-1^ to 0.42 *h*^-1^ in intervals of 0.02 *h*^-1^ and feeds with glucose concentrations of 5 g/l, 7.5 g/l, 10 g/l, 15 g/l and 30 g/l were simulated. In all cases the simulated cultures were glucose-limited, there was negligible glucose accumulation in the media, and the glucose mass balance was used to calculate the cell concentration in the reactor. If by-product secretion was predicted during simulations, their concentration was calculated using mass balances (Table 1). A function for chemostat simulations is provided in the supplementary material.

**Table 1.** Glucose, biomass and product mass balances in chemostat reactors. Chemostat reactors are in steady state and have a constant inflow and outflow of media at a rate F (l/h). Note that, the dilution rate, D, (h^-1^) is defined as F/V where V represents the reactor volume (l). The biomass balance shows that D equals the cell growth rate (*μ*, *h*^-1^). Given D and the glucose concentration in the feed (*c_s,in_*, mmol/l), FBA is used to calculate the glucose consumption rate (*q_s_*, mmol/*gDw*/h) and the glucose mass balance is used to calculate the cell concentration (*c_x_*, *gDw*/l). Similarly, the concentration of product i (*c_i_*, mmol/l) is calculated with the product mass balance using its production rate (*q_i_*, mmol/*gDw*/h) obtained by FBA.

Chemostat mass balances
|  |  |
| --- | --- |
| **Glucose** | $0 = D * c_{s,in} - q_s * c_x - D * c_s$ |
| **Biomass** | $0 = \mu * c_x - D * c_x$ |
| **Product i** | $0 = q_i * c_x - D * c_i$ |

#### 2.2.2. Batch and fed-batch simulations

During batch and fed-batch simulations the growth reaction (r_2111) was set as objective to maximize [18]. Following Sánchez et al., the upper bound of the protein pool reaction was increased by 25% [9]. When ethanol was present in the reactor, its uptake was allowed un-constraining reactions r_1761 (Yeast8) or r_1761_REV (ecYeast8).

During the simulation of batch and the batch phase of fed-batch reactors, the glucose exchange reaction (r_1714 or r_1714_REV) was constrained based on the glucose concentration in the reactor using a Michaelis-Menten kinetic equation (*q_glc,max_* = 10 mmol/*gDw*/h, *k_m,glc_* = 0.28 mmol/l [19]). The glucose mass balance was used to calculate the remaining glucose in the reactor. During the feeding phase of fed-batch reactors the glucose mass balance was used to calculate the glucose uptake rate and constrain the glucose exchange reaction. During this phase glucose is the limiting factor and its concentration in the reactor is negligible (Table 2).

**Table 2.** Glucose, biomass and product mass balances in batch and fed-batch reactors. In batch reactors and during the batch phase of fed-batch reactors biomass and products accumulate, substrates are depleted at a rate determined by the Michaelis-Menten equation and the glucose mass balance is used to calculate the remining glucose in the reactor (*M_s_*, mmol). During the feeding phase of fed-batch reactors, the reactors are fed with media containing substrates at a rate F (l/h), there is no accumulation of glucose (i.e. dMs/dt = 0) and the glucose mass balance is used to constrain the glucose uptake rate (*q_s_*, mmol/*gDw*/h). FBA is used to calculate the growth rate (*μ*, *h*^-1^) and the production rate of other metabolites (*q_i_*, mmol/*gDw*/h). Mass balances are used to calculate the biomass mass in the reactor (*M_x_*, *gDw*) as well as the product mass (*M_i_*, mmol). Note that mass is used instead of concentrations as the volume in the reactor changes during the feeding phase, concentrations are calculated as M/V(t), where V(t) is the liquid volume in the reactor at time (t).

Batch and fed-batch mass balances
|  |  |
| --- | --- |
| **Glucose** | Batch phase: $\frac{dM_s}{dt} = -q_s * M_x(t)$ |
| | Feeding phase: $0 = F * c_{s,in} - q_s * M_x$ |
| **Biomass** | $\frac{dM_x}{dt} = \mu * M_x(t)$ |
| **Product i** | $\frac{dM_i}{dt} = q_i(t) * M_x(t)$ |

After FBA optimization, the predicted metabolic fluxes were used to calculate new cell and by-products concentrations in the reactor using integrated mass balances (Table 2). The functions used for these simulations are available in the Supplementary Material.

Simulations of batch growth on combinations of sucrose and glucose, sucrose and fructose and sucrose and mannose were performed. During these simulations uptake of glucose, fructose, mannose and sucrose was allowed if these metabolites were present in the reactor by setting a negative lower bound (Yeast8) or a positive upper bound (ecYeast8) to their exchange reactions (r_1714 and r_1714_REV, r_1709 and r_1709_REV, r_1715 and r_1715_REV, r_2058 and r_2058_REV respectively). This upper bound was calculated using a Michaelis-Menten equation for glucose. For the rest of substrates a lower bound of −10 mmol/gDW/h was used in simulations with Yeast8 and the upper bound of these reactions was unbounded in ecYeast8. Supplementary simulations were performed with ecYeast8 including additional constraints based on inhibition of substrate uptake by some of the carbon sources (Appendix A.1). Scripts required for these simulations are available as Supplementary Material.

#### 2.2.3. *Simulation of* Δpdc *lactate producing* S. cerevisiae

Yeast8 and ecYeast8 were modified to simulate a strain without a pyruvate decarboxylase (PCD) activity and expressing the lactate dehydrogenase gene from *Lactobacillus plantarum* [20]. In both models the growth reaction (r_2111) upper bound was constrained to 0.13 *h*^-1^ to simulate the maximum growth rate observed experimentally [20]. dFBA was used to simulate cells growing in a 1 l reactor operated as a batch with 100 g/l of initial glucose and a 100 g glucose pulse 75 hours after inoculation. Oxygen limitation was experimentally observed 24 h after inoculation until the end of the process and was simulated constraining the oxygen exchange reaction (r_1992 in Yeast8 and r_1992_REV in ecYeast8) assuming pseudo-steady state for oxygen (i.e. no accumuation) [20]. During simulations with ecYeast8 the export of products different than biomass, lactate, succinate and glycerol was avoided constraining their secretion reactions [20]. Export of metabolites was unconstrained in simulations with Yeast8 to avoid infeasible solutions. A detailed explanation of model modifications and simulations can be found in Appendix A.2.

### 2.3. Experimental data

Experimental data of *S. cerevisiae* growth in chemostat and batch reactors was obtained from literature (Table 2).

Fed-batch cultures of *S. cerevisiae* CEN.PK-113-7D were performed in a 1 l working volume of a stirred fermenter (DASGIP parallel bioreactor system, Eppendorf). Throughout the fermentations the pH was kept at 5.1 and the temperature was kept at 30 degrees Celsius. The batch medium (400g) contained: 2.5 g/kg glucose, 1g/kg (NH)4SO4, 10g/kg KH2PO4, 4g/kg MgSO4*7H2O, 0.3g/kg CaCl2*2H2O and vitamins and trace elements according to Verduyn et al. [21]. After 4 hours of growth on the batch medium the aerobic fed-batch phase was started with an exponential feed profile supporting a growth rate of 0.05 *h*^-1^. The composition of the feed medium was 209 g/kg glucose, 7.67g/kg ethanol, 2g/kg (NH)4SO4 20g/kg KH2PO4, 8g/kg MgSO4*7H2O, 0.6g/kg CaCl2*2H2O and vitamins and trace elements according to Verduyn et al. [21]. Samples for biomass concentration determination were obtained every 24 hours and actual oxygen uptake rate and CO2 production rate were determined using off-gas analysis.

All experimental data is available as Supplementary Data.

**Table 2:** Summary of experimental data used in this study. *CEN.PK 113.7D pdc1(−6.−2)::loxP pdc5(−6.−2)::loxP pdc6(−6.−2)::lox P ura3-52 YEpLpILDH

| Reactor operation | Carbon source | Strain | Reference |
| --- | --- | --- | --- |
| Chemostat | Glucose | CBS8066 | [22] |
| Chemostat | Glucose | DS28911 | [23] |
| Chemostat | Glucose | H1022 | [24] |
| Chemostat | Glucose | CEN.PK 113.7D | [25] |
| Batch | Glucose | H1022 | [26] |
| Fed-batch | Glucose | CEN.PK 113.7D | This study |
| Batch | Sucrose + Glucose | DGI342 | [27] |
| Batch | Sucrose + Fructose | DGI342 | [27] |
| Batch | Sucrose + Mannose | DGI342 | [27] |
| Batch | Glucose | GCSI-L* | [20] |

### 2.4. Sampling of intracellular fluxes

The Artificial Centering Hit-and-Run (ACHR) Sampler module from cobrapy (achr) was used to sample the solution space. Before sampling, upper and lower bounds of the biomass reaction were constrained to the desired growth rate, glucose uptake was set as objective to minimize and the flux through this reaction was constrained to the minimal flux ± 10%. In all cases the modified model was used, 10000 samples were taken and their feasibility, understood as the satisfaction of mass balances and maintenance constraints, was validated using achr.validate function. Samples were taken at a range of growth rates (0.01, 0.05, 0.1, 0.15, 0.2, 0.23, 0.24, 0.25, 0.26, 0.27, 0.28, 0.29 and 0.3 *h*^-1^). To analyze intracellular fluxes the median ± the median absolute deviation (MAD) of the valid samples is considered as the predicted flux through a given reaction. Sampling data is presented as Supplementary Data.

## 3. Results

We used FBA to simulate *S. cerevisiae* chemostat growth and dFBA to simulate batch and fed-batch runs. We compared predictions of Yeast8 and ecYeast8 with experimental data. In all cases, the initial conditions in the reactor were used as inputs for the simulation. FBA and mass balances were used to calculate metabolic fluxes and concentrations in the reactor. Besides, flux sampling of ecYeast8 was used to predict active central carbon metabolic fluxes at different growth rates.

### 3.1. Chemostat simulations

*S. cerevisiae* cells grown in continuous cultures change their metabolism depending on the dilution rate. At low growth rates they present a completely aerobic metabolism whereas ethanol production is observed at growth rates higher than the critical dilution rate (*D_crit_*), process known as the Crabtree effect. Data from chemostat growth of *S. cerevisiae* strains CBS8066, DS28911 and H1022 was obtained from [22]–[24]. In these experiments *S. cerevisiae* was grown at different dilution rates (*D*) with different glucose concentrations in the feed. For each dilution rate, we constrained the growth rate of Yeast8 and ecYeast8 and used mass balances to calculate the cell, glucose and by-product concentrations in the reactors at steady state.

Figure 1 A shows predictions of biomass concentrations by Yeast8 and ecYeast8. Whereas Yeast8 predicts constant biomass concentration, ecYeast8 simulates a decrease in biomass concentration after a specific dilution rate, the critical dilution rate, *D_crit_*. The decrease in biomass concentration is also observed in the experimental data, which shows different critical dilution rates for different strains [28]. The model predicts a critical dilution rate of 0.27 *h*^-1^, in agreement with that reported for strains DS28911 and H1022 (0.28 *h*^-1^ and 0.21 *h*^-1^) [23], [24]. Strain CBS8066 has a higher protein content than H1022 and shows a higher critical dilution rate (0.38 *h*^-1^)[22], [29]. This higher growth rate was simulated increasing protein availability in the model (26.8% increase of the upper bound of the protein pool reaction) showing that tuning protein availability results in different *D_crit_*, suitable to predict chemostat growth of different *S. cerevisiae* strains.

**Figure 1.**
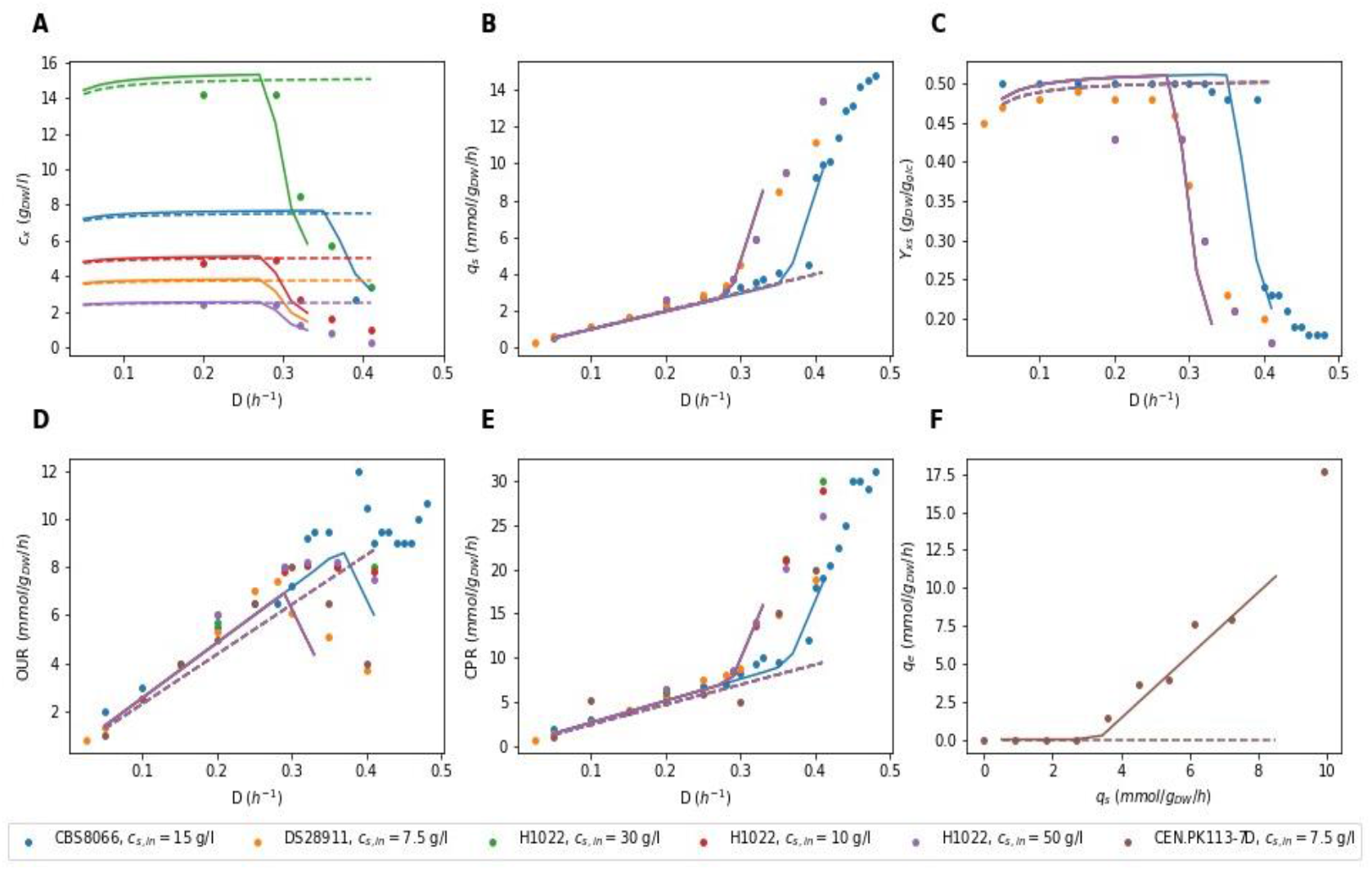
Chemostat simulations with Yeast8 (- -) and ecYeast8 (-) compared with experimental data. (A) Biomass concentration (*c_x_*), (B) Specific glucose uptake rate (*q_s_*), (C) Yield of biomass on glucose (*Y_xs_*), (D) Specific oxygen uptake rate (OUR) and (D) Specific *CO*_2_ production rate (CPR) at different dilution rates (*D*). (F) Specific ethanol production rate (*q_e_*) at different specific glucose uptake rates (*q_s_*). Experimental data of strains CBS8066, DS28911, H1022 and CEN.PK 113.7D where obtained from [22]–[25]respectively. Note than in figures B-F all dashed lines overlap and continuous orange, green and red lines overlap with the continuous purple line.

Figure 1 also shows the maximum growth rate predicted by the model with the default bound for the protein pool reaction is 0.30 *h*^-1^ (0.38 *h*^-1^ when this bound is increased) while all strains are able to grow at dilution rates as high as 0.4 *h*^-1^. However, when cells are grown experimentally at dilution rates higher than 0.3 *h*^-1^, the dilution rate has to increase in small steps to avoid wash out, indicating cells need time to adapt to high growth rates [24]. This adaptation is related with an increase in protein content and therefore, chemostat predictions at high growth rates would improve with a growth rate dependent protein availability constraint. Interestingly, decreasing maintenance requirements in the model did not affect maximum growth rates predictions, suggesting that the protein availability constraint implicitly accounts for protein synthesis costs and reduces the impact of the maintenance reaction in the simulations.

According to simulations with Yeast8, specific glucose uptake is proportional to the dilution rate. However, experimental data and simulations with ecYeast8 show a sharp increase in glucose uptake after *D_crit_* (Figure 1 B). Higher glucose uptake rates and lower biomass concentrations result in a decrease on the biomass yield on glucose after *D_crit_*, which is only predicted by simulations using ecYeast8 (Figure 1 C).

Similar to the glucose uptake rate predictions by Yeast8, oxygen uptake rates and *CO*_2_ production rates are predicted to be proportional to the growth rate (Figures 1 D and E). However, after *D_crit_* cells show a partially fermentative metabolism that results in a decrease of the oxygen uptake rate and an increase on the *CO*_2_ production rate. Besides, ecYeast8 predicts byproduct formation at growth rates higher than the critical dilution rate. It predicts secretion of acetaldehyde and acetate and accurately predicts ethanol flux at different glucose uptake rates (Figure 1 F). None of these changes are predicted by Yeast8.

### 3.2. Batch and fed-batch simulations

During batch fermentations glucose is present in excess and cells grow at their maximum growth rate. Yeast8 and ecYeast8 were used to simulate batch growth of *S. cerevisiae* and the results were compared to experimental data [26]. Glucose uptake rate was constrained in both models as a function of the glucose concentration in the reactor according to Michaelis-Menten kinetics. Whilst glucose uptake was the only constraint imposed to Yeast8, ecYeast8 was also limited by the availability of proteins.

Simulations using Yeast8 predict no ethanol production, faster glucose consumption and higher cell concentrations than the experimental measurements (Figure 2). In these simulations glucose uptake kinetics determines how fast glucose is consumed and all fluxes are distributed to optimize biomass production which results in exponential growth, no by-product formation as well as glucose depletion and growth arrest after 5h.

**Figure 2.**
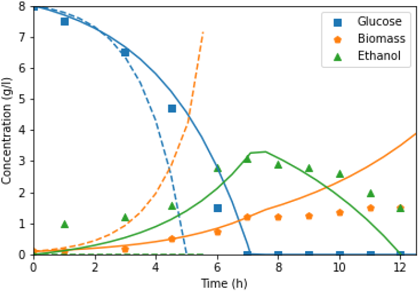
dFBA simulation of batch growth of *S. cerevisiae* H1022 with Yeast8 (- -) and ecYeast8 (-) compared with experimental data (symbols) [26].

Contrarily, simulations using ecYeast8 accurately predict glucose and biomass concentrations until glucose is depleted 7h after inoculation. This model also predicts the production of ethanol and its consumption after glucose depletion. During these simulations the growth rate is limited by protein availability, and only at glucose concentrations approaching *k_m_* (0.28 mmol/l), the Michaelis-Menten equation for glucose uptake becomes the limiting factor. The protein availability constraint results in ethanol production by ecYeast8 and a realistic yield of biomass on glucose, overestimated by Yeast8. Although ethanol consumption was allowed during the entire simulation, it was only predicted after glucose depletion (in agreement with experimental data). However, during this phase ecYeast8 simulates higher biomass concentration and faster ethanol consumption than the experimental measurements.

In fed-batch reactors, batch growth is followed by a feeding phase in which media with substrate enters the reactor. During this phase cellular growth is determined by the available glucose. We performed fed-batch cultivation of *S. cerevisiae* CEN PK-113-7D and used oxygen uptake and *CO*_2_ production rates (OUR, CPR) as an indication of cell metabolism. This process was simulated using dFBA and model predictions were compared to experimental data. Yeast8 showed higher OUR and CPR as a result of a higher growth rate during the batch phase which resulted in depletion of glucose before the start of the feeding phase. Simulations with ecYeast8 resulted in accurate prediction of OUR, CPR and biomass concentration in the reactor (Supplementary Figure B.6).

### 3.3. Batch growth on multiple carbon sources

Yeast8 and ecYeast8 were used to simulate batch growth of *S. cerevisiae* in a mixture of carbon sources using dFBA. Dynesen et al. [27] combined sucrose, a disaccharide of glucose and fructose, with glucose, fructose or mannose in order to study growth and catabolite repression of *S. cerevisiae* DGI342. Simulations using Yeast8 and ecYeast8 were compared with this experimental data.

Yeast8 predicts simultaneous consumption of all carbon sources and unrealistically high uptake rates resulting in substrate depletion after 6 hours (Figure 3 A-C). In order to obtain better predictions, uptake reactions should be constrained using specific Michaelis-Menten kinetic equations for each carbon source.

**Figure 3:**
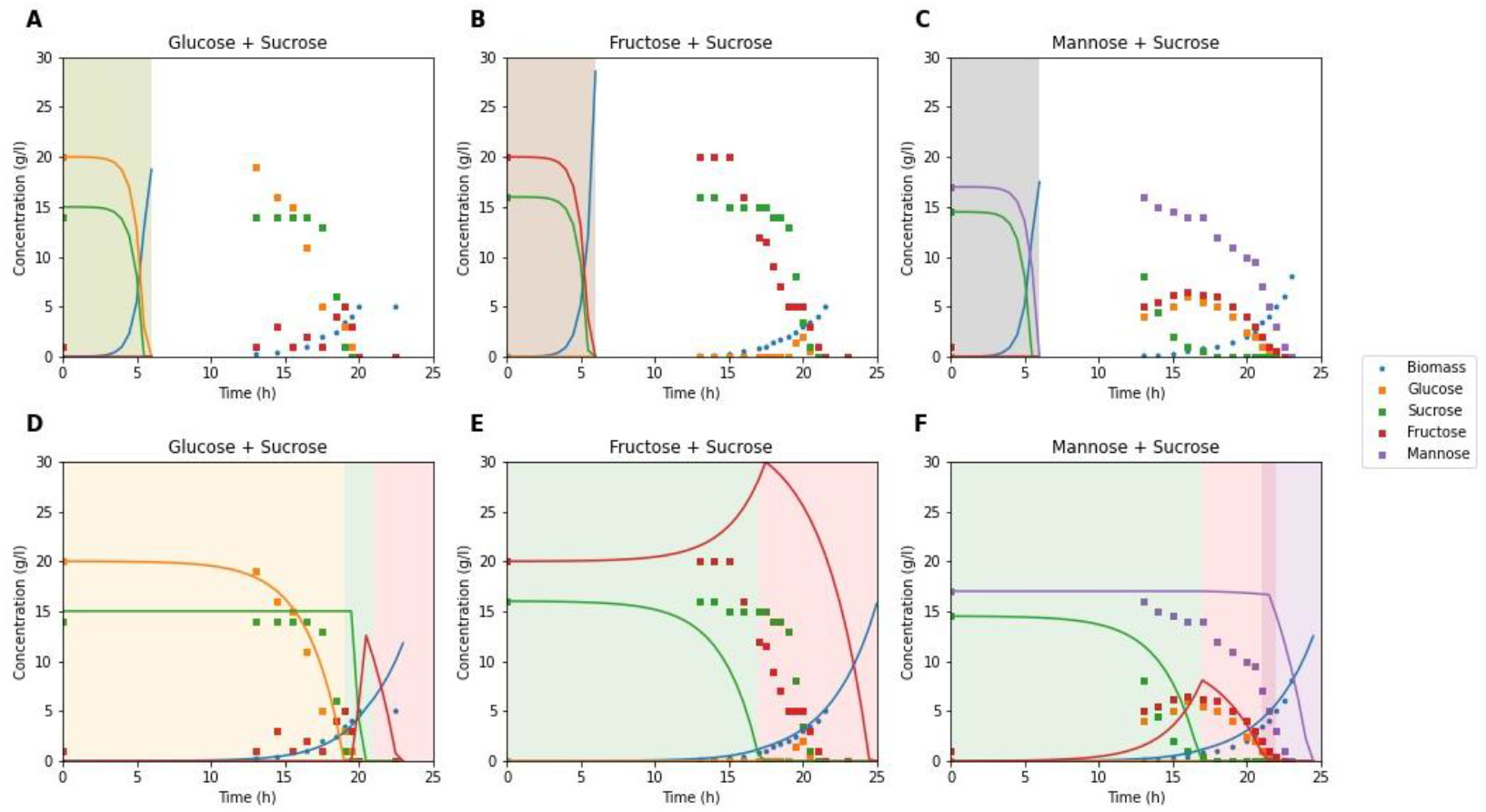
Two-carbon sources *S. cerevisiae* DGI342 batch simulations using Yeast 8 (A, B, C) and ecYeast8 (D, E, F) compared to experimental data (symbols) [27]. Colored areas represent different substrate consumption phases predicted by the model: glucose consumption (orange), sucrose hydrolysis and glucose consumption (green), fructose consumption (red) and mannose consumption (purple).

Contrarily, ecYeast8 simulations show a good agreement with experimental data as the order of substrate consumption in this model is determined by the relative protein cost for substrate consumption as well as the biomass yield on the different carbon sources (Table 3) (Figure 3 D-F).

**Table 3.** Relative protein cost for consumption of different substrates and biomass yield per C-mol. The relative protein cost is calculated as the flux through the protein pool reaction required to consume 1 mmol of substrate divided by the same flux required for consumption of 1 mmol of glucose.

| Substrate | Relative protein cost | Biomass yield [gDW/C-mol] |
| --- | --- | --- |
| Glucose | 1 | 0.43 |
| Fructose | 1.25 | 0.30 |
| Mannose | 1.27 | 0.30 |
| Sucrose | 2.18 | 0.20 |

When sucrose and glucose are the substrates, the model predicts three phases characterized by the use of different carbon sources. First, all the available free glucose is consumed, as it is the substrate with the lowest protein cost (Table 3). In the second phase sucrose is hydrolyzed, sucrose-derived glucose is consumed and fructose accumulates. The highest protein cost of sucrose is caused by the simultaneous consumption of glucose and fructose. However, during dFBA simulation the accumulation of glucose and fructose in the reactor is allowed and the only additional cost of sucrose consumption is caused by the need to hydrolyze the disaccharide by the invertase enzyme. After hydrolysis, ecYeast8 predicts glucose consumption and fructose accumulation due to the lower protein cost of glucose degradation (Table 3). The third phase is characterized by fructose consumption, with a higher protein cost compared to glucose caused by a higher flux through the glucose-6-phosphate (G6P) isomerase. According to ecYeast8, this enzyme converts G6P to frucotse-6-phosphate (F6P) during growth on glucose and catalyzes the reversible reaction during fructose growth with a higher flux. The fact that these three phases are also observed experimentally suggests that carbon catabolite repression (CCR) of sucrose, fructose and ethanol exerted by glucose is essential to achieve maximum growth rate when considering the limitation of protein content in the cells (Figure 3 D).

When the carbon sources are sucrose and fructose, the model repeats phases two and three. First, sucrose is hydrolyzed, the sucrose-derived glucose is consumed and fructose accumulates. Then, only after glucose depletion, the model consumes the available fructose (Figure 3 E).

Simulations with sucrose and mannose show similar results. First, the model predicts consumption of sucrose-derived glucose and fructose accumulation. Fructose consumption only starts after glucose depletion. Mannose consumption starts at a low rate at the end of the fructose consumption phase and continues then at a higher rate due to the higher protein cost required for its degradation. This cost is caused by the need to convert mannose to F6P, reactions catalyzed by mannokinase and mannose-6-phosphate isomerase (Figure 3 F).

Although the model does not predict initial consumption of fructose in simulations with fructose and sucrose or initial glucose accumulation and simultaneous consumption of glucose, fructose and mannose in sucrose and mannose simulations, the protein availability constraint is enough to accurately predict sucrose hydrolysis as well as fructose and mannose consumption rates. Besides, the combination of ecYeast8 and dFBA improved predictions by explicitly modelling the inhibitory effect of glucose, fructose, sucrose and mannose on the uptake rates of the other carbon sources (Supplementary Figure B.7).

### 3.4. *Simulation of* Δpdc *lactate producing* S. cerevisiae

Yeast8 and ecYeast8 were modified to simulate a *S. cerevisiae* strain without pyruvate decarboxylase activity, laboratory evolved to tolerate high glucose concentrations and engineered to produce lactate[20]. dFBA simulations with Yeast8 and ecYeast8 were in agreement with experimental data (Figure 4A). According to van Maris et al. [20] during the first 24 h of the fermentation oxygen was supplied in excess to the reactor and cells were only limited by glucose availability. After 24 h cells suffered oxygen limitation, which was simulated constraining the oxygen uptake reaction. The oxygen limitation continued after 75 hours when additional 100 g of glucose were added to the reactor.

**Figure 4.**
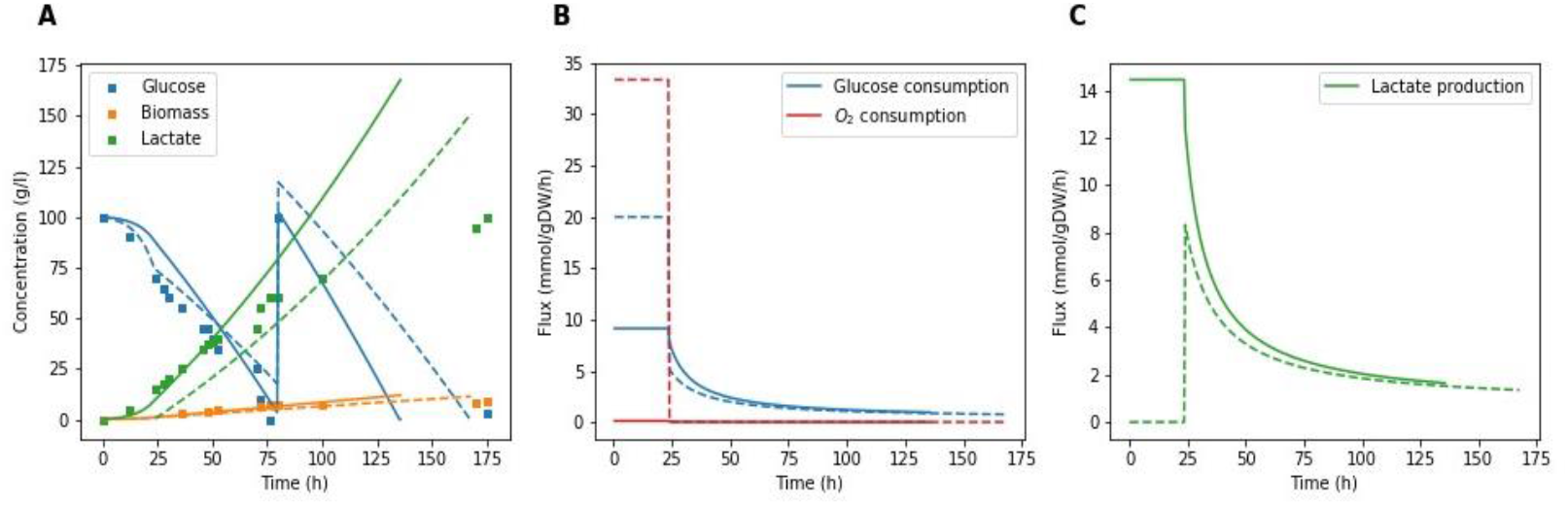
Batch growth simulation of Δ*pdc*, lactate producing *S. cerevisiae* using EcYeast8 (-) and Yeast8 (- -) compared to experimental data [20].

During laboratory evolution fastest growers were selected, obtaining a final strain with a maximum growth rate of 0.13 *h*^-1^ [20]. Although the concept of laboratory evolution is in agreement with the use of biomass growth as objective function during FBA, Yeast8 and ecYeast8 predicted higher maximum growth rates, which suggests that the obtained strains could be further evolved. Therefore, the upper bound of the biomass reactions had to be constrained to match the experimental value.

During the glucose limitation phase both models, Yeast8 and ecYeast8, were limited by glucose availability determined by a Michaelis-Menten equation. Besides ecYeast8 was limited by the protein pool constraint which resulted in prediction of a lower glucose uptake rate by this model (Figure 4 B). In this period oxygen uptake rates predicted by Yeast8 where unreasonably high and lactate production was not predicted (Figure 4 B, C). Contrarily, the limitation in protein availability of ecYeast8 resulted in realistic predictions of oxygen uptake and lactate production rates. After 24 h the limitation in oxygen uptake resulted in a 99.88% decrease in oxygen uptake by Yeast8 (from 34 mmol/*gDw*/h to 0.04 mmol/*gDw*/h) and 93% decrease in ecYeast8 (from 0.58 to 0.04 mmol/*gDw*/h) (Figure 4). After the introduction of this limitation there were not significant differences in flux predictions by both models.

Besides lactate production, simulations by ecYeast8 resulted in succinate and glycerol production at concentrations similar to experimental measurements [20]. Simulations with Yeast8 only resulted in glycerol and succinate production once oxygen uptake was limited and additional by-products such as citrate or arginine were exported by the model.

Although both models performed well when the observed oxygenl imitation was included in the model, the protein availability constraint was enough to predict lactate production in oxygen excess conditions suggesting the potential of combining enzyme constrained models and dFBA for cell factory simulations (Figure 4). The disagreement between model predictions and experimental data observed during the second half of the simulations is probably caused by growth inhibition due to product toxicity. The dFBA framework would allow to include this inhibition linking the upper bound of the biomass reaction to the reactor concentration of the toxic compound.

### 3.5. *Flux sampling as tool to explore* S. cerevisiae *metabolism at different growth rates*

When modelling cell metabolism, FBA only provides one of the multiple flux distributions that results in the optimization of the chosen objective function. Flux sampling algorithms solve this problem by providing possible flux distributions of metabolic reactions that satisfy mass balance constraints [15]. Due to the better performance of ecYeast8 when simulating consumption and production of metabolites in a reactor setting, we tested how flux sampling can be applied to study intracellular fluxes.

During simulations with ecYeast8, all the glyceraldehyde-3-phosphate was produced through the pentose phosphate pathway. To ensure experimentally observed flux though phospho-fructo kinase and fructose bisphosphate aldolase, the reversible transaldolase reaction was blocked before sampling [30]. According to Pereira et al. [31] reversibility of cytosolic reactions should favor the production of NADH. Therefore, the reduction of tricarboxylic acid cycle (TCA) intermediates in the cytoplasm was avoided. Lastly, the model was re-scaled to avoid stoichiometric coefficients below solver tolerance that caused numerical instability.

For each simulation the obtained flux distributions represents the metabolism of *S. cerevisiae* cells growing in a chemostat with a specific dilution rate at steady state. Besides, the simulation at maximum growth rate represents the metabolism of *S. cerevisiae* cells growing exponentially in a batch reactor. Sampling results can be found as supplementary data.

In general, we observed good agreement between predicted fluxes and experimental measurements (Supplementary Figure B.8). As example, the flux through TCA reactions decrease with growth rate and, at maximum growth rate, sampling results show the operation of the TCA cycle as two different branches (zero flux through *α*-ketoglutarate dehydrogenase (KGD), succinyl-CoA synthetase (SCL) and fumarase (FUM))[30], [32], [33]. At these high growth rates relative flux to the pentose phosphate pathway (PPP) decreases and is directed towards glycolysis and ethanol formation (Figure 5A). As expected, the variability of the fluxes decreases at increasing growth rates as a result of a more limited solution space. At higher growth rates protein availability becomes limiting and alternative pathways are no longer feasible.

**Figure 5.**
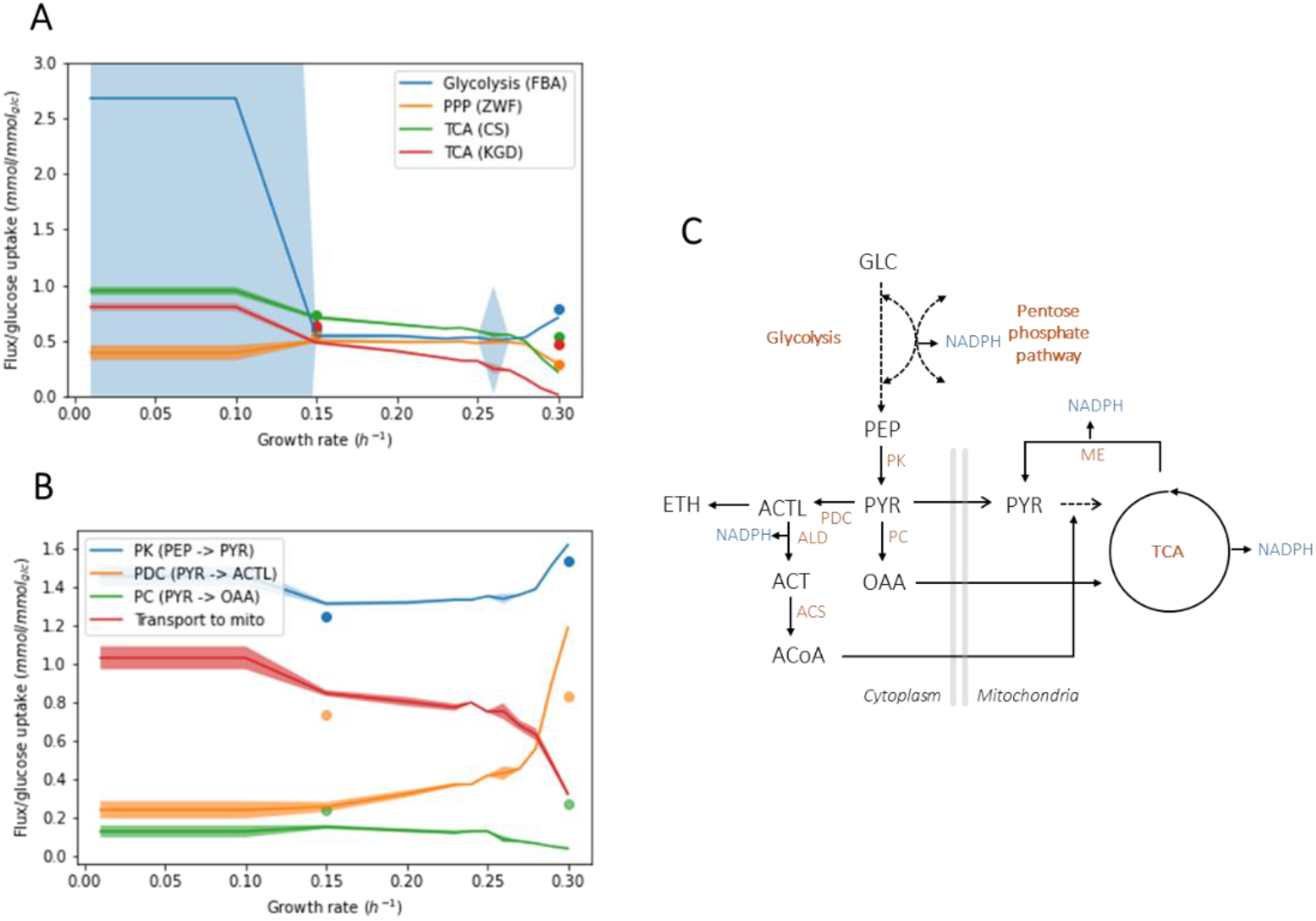
Comparison of flux sampling results with ecYeast8 (median ± MAD) and 13C flux analysis data (symbols) [30]. (A) Fluxes relative to the glucose uptake at different growth rates of the glycolytic enzyme fructose bis-phosphate aldolase (FBA), the pentose phosphate pathway enzyme glucose-6-phoaphate dehydrogenase (ZWF), the tricarboxilic acid cycle enzymes citrate synthase (CS) and alpha-keto glutarate dehydrogenase (KGD). (B) Fluxes relative to the glucose uptake of enzymes involved in the pyruvate node (pyruvate kinase, PYK; pyruvate decarboxylse, PDC; pyruvate carboxylase, PC) and relative transport flux of pyruvate to mitochondria (mito). (C) Representation of the pyruvate node (GLC, glucose; PEP, phosphoenol pyruvate; PYR, pyruvate; OAA, oxaloacetate; ACTL, acetaldehyde; ETH, ethanol; ACT, acetate; ACoA, acetyl coenzyme A; ALD, acetaldehyde dehydrogenase; ACS, acetyl CoA synthase; ME, malic enzyme).

As test case on the use of flux sampling to study metabolism, we focused on the predicted flux distributions in the pyruvate node and compared them to experimental data (Figure 5B, C) [30]. Pyruvate kinase (PK) is the main source of cytoplasmic pyruvate and, in agreement with literature, the model predicts constant relative flux at growth rates below the critical dilution rate and increasing relative flux at higher growth rates. Whilst flux predictions of PK and pyruvate decarboxylase (PDC) follow the same trend as experimental data, pyruvate carboxylase (PC) shows the opposite behavior (Supplementary Figure B.8). Frick and Wittmann [30] propose that at high growth rates pyruvate conversion to acetyl-CoA (by PDC, ALD and ACS) and subsequent transport to mitochondria saturates. The extra pyruvate is then converted to oxaloacetate by PC, which is transported to the mitochondria and converted back to pyruvate by the malic enzyme (ME). In this way the mitochondrial pyruvate pool, required for acetyl-CoA and amino acid synthesis, is replenished [30], [34]. However, the model predicts a relative flux increase through PDC, no saturation in the cytoplasmic conversion of pyruvate to acetyl CoA and, as a result, fails to predict the experimentally observed flux increase through PC (Figure 5 C). Although model predictions show a decrease in relative pyruvate transport to the mitochondria, transport is enough to cover mitochondrial pyruvate requirements and the experimentally observed flux increase through ME is not predicted by the model (Figure 5 C). In fact, free movement of metabolites across compartments is allowed in ecYeast8 as transporters are not part of the protein pool. Therefore, inaccurate flux predictions are expected when transport of metabolites across compartments is the limiting factor.

## 4. Discussion

EcModels add an additional layer of information to traditional GEMs based on the limited capacity of the cells to synthesize proteins. We show here that this results in more accurate predictions of extracellular fluxes during chemostat, batch and fed-batch growth of different *S. cerevisiae* strains.

De Groot et al. [35] show that GEM predict overflow metabolism when two growth-limiting constraints are hit regardless of their biological interpretation. Therefore, Yeast8 can be modified to predict overflow metabolism by adding a second constraint such as a maximum oxygen uptake rate [36]. However, ecYeast8 not only predicts respiro-fermentative metabolism at growth rates higher than the critical dilution rate but, when combined with dFBA, it also accurately describes ethanol production and consumption during exponential growth, the preferred consumption order of different carbon sources as well as product production rates (Figures 2, 3, 4). In traditional GEM, the flux through reactions required for growth is not constrained, so the model adjusts these fluxes to obtain the desired growth rate, which results in inaccurate description of metabolism EcYeast8 breaks the linear dependency between fluxes and growth rate and shows accurate intracellular flux predictions (Figure B.8). Simulating this behavior with Yeast8 is only possible upon an iterative, case-dependent design of condition-specific constraints [37].

The parameter with the largest influence on the simulations is the upper bound of the protein exchange reaction, which represents enzyme availability in the model. This parameter determines the maximum growth rate in batch reactors, the critical dilution rate in continuous cultures and the uptake rates of different substrates. In the absence of proteomic data Sánchez et al. [9] assume constant protein availability for a given strain and process and provide two different values depending on the simulation of chemostat or batch growth. We showed that increasing this parameter was required to simulate batch growth on different carbon sources and that it should be adjusted to accurately simulate chemostat growth of strains with different protein content [29]. In fact, the protein contents of a specific strain in a given process should be tuned considering the expected glucose uptake and/or growth rate[1], [23], [30], [38]. Interestingly, the effect of the protein availability constraint is so strong that it implicitly accounts for protein synthesis costs reducing the impact of the maintenance requirements during the simulations. This suggests that the constraint in protein availability can be understood in terms of the limited space in the cell, but also in terms of limited energy available for protein synthesis. Besides, simulations with different carbon sources suggested that the order of substrate consumption can be partially explained by the associated protein cost required for its consumption. When considering the limitation of protein content in the cells, CCR is essential to achieve the maximum growth rate.

dFBA is a valuable tool to predict the dynamic behavior of engineered strains in a reactor context [12]–[14]. Whilst FBA allows the comparison of yields between different engineered strains, dFBA simulates dynamic processes allowing the comparison of final titters and productivities which depend, not only on the strain, but also on the bio-process. We showed here how the combination of ecYeast8 with (d)FBA improved predicted metabolic changes in response to the operation of a reactor without the need of additional constraints. We used simulations on mixture of carbon sources to show how predictions can be further improved incorporating regulation-related constraints to the dFBA framework (Supplementary Figure B.7). In this way, substrate uptake, growth rate and/or known metabolic reactions fluxes can be limited based on metabolite concentrations in the media [38]. We showed the potential of this method to aid the design of bio-processes including the prediction of the metabolism of engineered cells in a reactor and changes in cell metabolism due to co-feeds, a novel trend in bio-process design [39]. Finally, the effect of other process parameters such as temperature could also be included in the dFBA framework using temperature constrained ecModels [40].

Accurate predictions of intracellular metabolic fluxes is a desired feature for models aiming to find and compare metabolic engineering strategies to improve production of a target metabolite. To the best of our knowledge, this study is the first report on how to combine flux sampling and ecModels to study intracellular flux predictions, avoiding the necessity to fix an objective function and allowing the coverage of the whole solution space [15]. While previous studies focused on the prediction of intracellular fluxes at maximum growth rate, we have compared flux predictions covering *S. cerevisiae* full range of growth rates [31]. We showed that the protein availability constraint breaks the linear dependency between fluxes and growth rate and results in accurate intracellular flux predictions (Figure 5). As a test case, this strategy allowed us to find flux inconsistencies in model predictions of the pyruvate node, a key point in central metabolism (Figure 5). Despite of the substantially improved predictive power of the model, however the protein availability constraint was not enough to yield accurate predictions of all intracellular fluxes due to the highly dimensional solution space and the absence of regulatory information in the model. Predictions could improve considering space limitation in cell membranes as additional constraint or by the creation of an ensemble model [41], [42]. Ultimately, when ecModels are applied to strain design, this limitation can be overcome using proteomic data instead of a single constraint on the protein content of the cells[9]. Better intracellular flux predictions by ecModels will likely result in a reduction of the prediction of incorrect knock-out and overexpression targets when these models are used to improve the performance of cell factories.

## 5. Conclusions

We introduced flux sampling as a tool to analyze intracellular flux predictions of ecModels, of major importance for model guided strain design. We have shown how the successful combination of ecModels and (d)FBA allows the prediction of interactions among cell metabolism and the environment allowing the simulation of different strains under different (and dynamic) production conditions. As parameters in the reactor as well as genetic modifications affect flux predictions, the combination of ecModels and dFBA allows the comparison of yields and productivities among different strains and (dynamic) production processes. This model and simulation framework, which links intracellular processes to bioreactor operations therefore provides the means for more accurate and realistic designs of cell-based processes increasing their usefulness for industrial applications.

## 7. Acknowledgment

This project was founded by NWO (project number GSGT.2019.008) and IBISBA (H2020 project numbers 730976 and 871118).

## 8. Author contributions

Conceptualization: SMP/JS/VAPMdS/MSD; Data curation:SMP; Formal analysis:SMP/MSD; Funding acquisition:VAPMdS; Methodology; SMP/ JS/VAPMdS/MSD Project administration VAPdMS/MSD; Supervision:JS/ VAPMdS/MSD; Visualization: SMP; Roles/Writing - original draft: SMP; Writing - review & editing: SMP/JS/VAPMdS/MSD.

## Appendix A. Supplementary methods

### Appendix A.1. Simulation of mixed carbon fermentations including additional regulation with ecYeast8

Consumption of combinations of sucrose with glucose, fructose and mannose were simulated using ecYeast8 and including additional regulatory constraints in the dFBA framework.

First, repression of fructose consumption by glucose was simulated by constraining the upper bound of the fructose uptake reaction (r_1709_REV) to zero when glucose concentration in the media was higher than 1 mmol/l. Similarly, repression of glucose consumption by fructose was simulated constraining the glucose uptake and transport reactions (r_1714 REV and r_1166) to zero when fructose concentration in the media was above 1 mmol/l. When sucrose and mannose were the initial carbon sources, the experimentally observed delay between sucrose hydrolysis and glucose and fructose consumption was simulated constraining their transport reactions (r)1166 and r_1134) to zero when sucrose was present in the media.

Second, sucrose hydrolysis was forced when glucose and fructose concentrations in the media were below 15 g/l by setting a lower bound to reaction r_2058 REV. If glucose and fructose were initially present in the media the bound was 3.5 mmol/*gDw*/h and 20 mmol/*gDw*/h otherwise.

In all cases, lower and upper bounds of glucose, fructose and mannose uptake as well as sucrose hydrolysis were set to zero when the carbon sources were not present in the media.

Last, during simulations with fructose and mannose as initial carbon sources, the protein exchange upper bound was increased by 75% compared to batch simulations with glucose and the secretion reactions for ethanol, acetate, pyruvate, acetaldehyde and 2,3-butanediol were unconstrained.

### Appendix A.2. *Simulation of* Δpdc *lactate producing strain*

Yeast8 and ecYeast8 were modified to simulate a strain without a pyruvate decarboxylase (PDC) activity and expressing the lactate dehydrogenase gene from *Lactobacillus plantarum* [20]. To simulate the knock out, bounds of reactions r_0959 and r_0960 (Yeast8) and arm r_0959 and arm r_0960 (ecYeast8) were set to zero. The lactate dehydrogenase reaction was added to Yeast8 (Equation A.1). Three reactions were added to ecYeast8: the forward and reverse reactions of the lactate dehydrogenase (Equations A.2, A.3) and a draw reaction for the LDH protein (Equation A.4). Values of *k_cat_* (40 *s*^-1^) and molecular weight (39 kDa) were obtained from BRENDA [43].

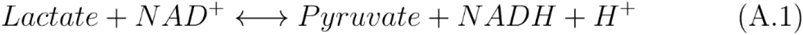

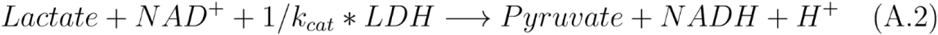

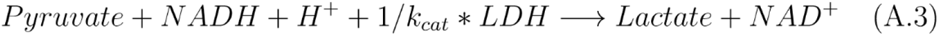

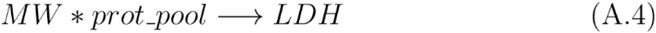

Batch growth of both model was simulated using dFBA. The growth reaction (r_2111) upper bound was constrained to 0.13 *h*^-1^ to simulate the maximum growth rate observed experimentally [20]. Cells were simulated in a 1 l reactor operated as a batch with 100 g/l of initial glucose. Eighty hours after inoculation 100 g of glucose were added to the reactor and this pulse was included in the simulations. According to [20] cells experienced oxygen limitation from 24 h after inoculation until the end of the process. Oxygen limitation was simulated constraining the oxygen uptake reaction (r_1992 in Yeast8 and r_1992_REV in ecYeast8) to *q_o_* assuming pseudo-steady state for oxygen:

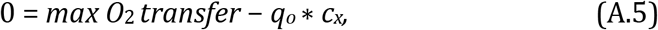

where the maximum *O*_2_ transfer to the reactor (*max O*_2__*transfer*) was calculated based on experimental data and *c_x_* is the predicted biomass concentration in the previous time step [20].

In agreement with experimental data, during ecYeas8 simulations export of products different than biomass, lactate, succinate and glycerol was avoided constraining their production reactions [20]. The same approach resulted in infeasible solutions with Yeast8 and therefore production of alternative products was allowed.

## Appendix B. Supplementary figures

**Figure B.6:**
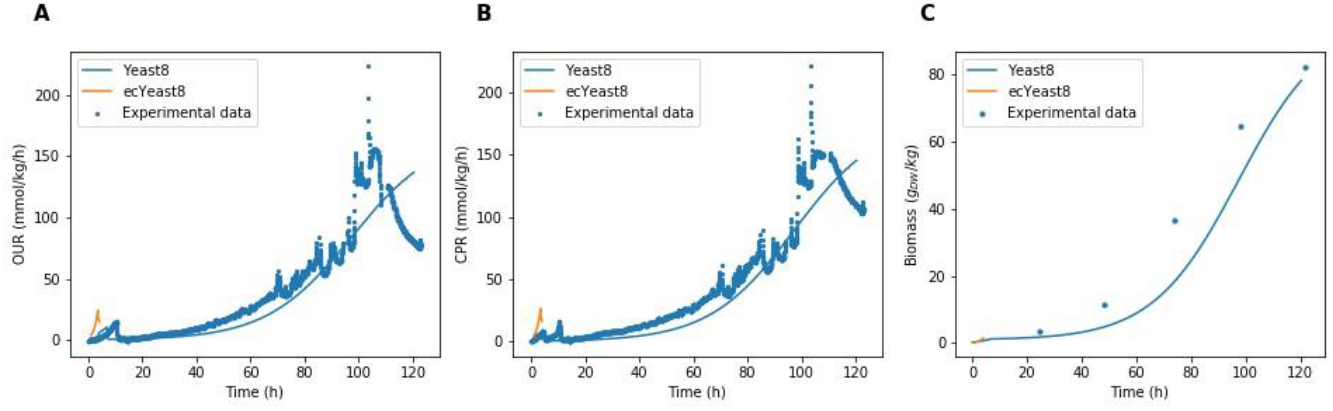
Simulations of OUR (A), CPR(B) and biomass concentration (C) of *S. cerevisiae* CEN.PK 113.7D grown with an exponential glucose feed. Peaks in experimental OUR and CPR data were caused by sampling of the reactor.

**Figure B.7:**
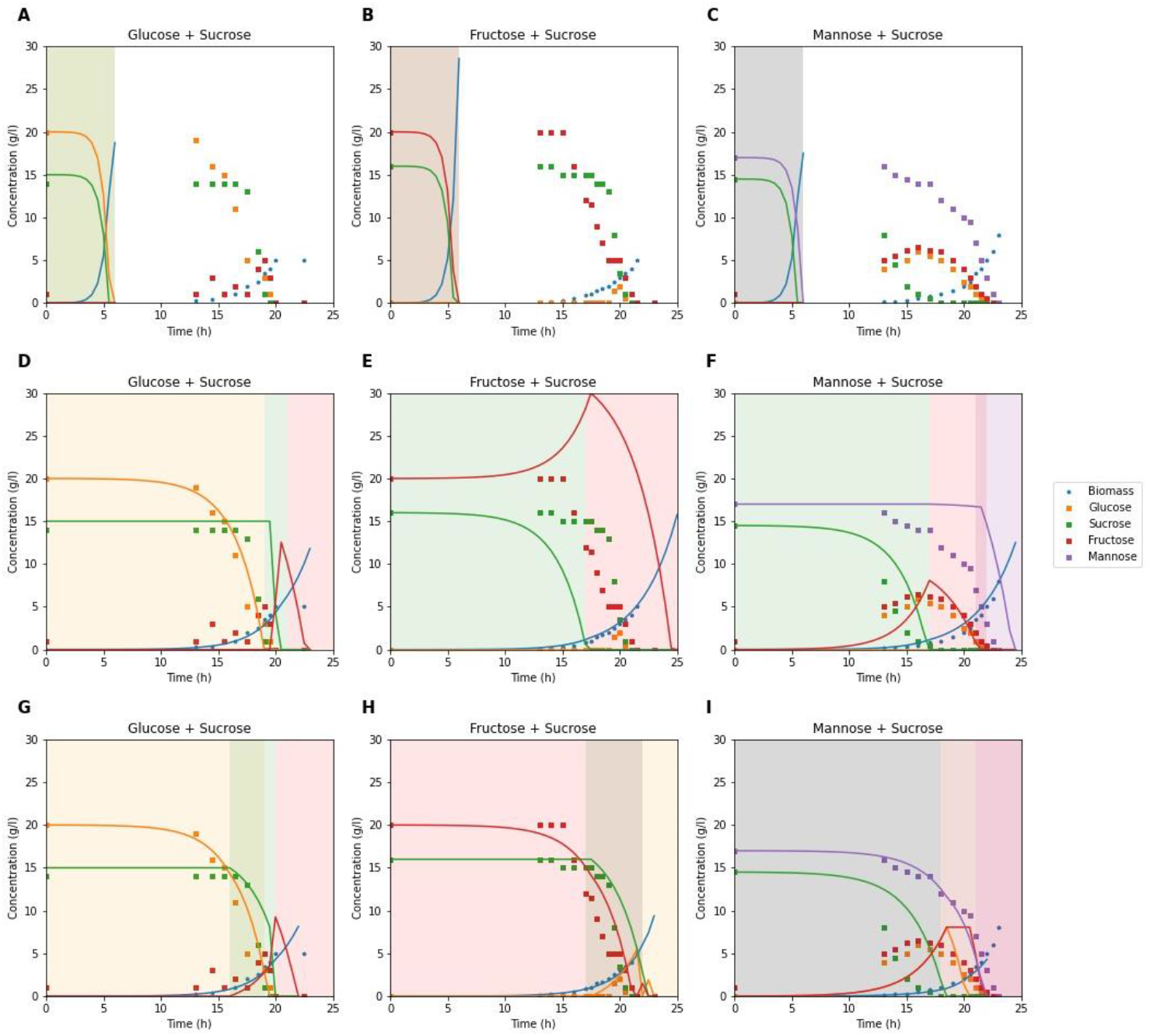
Two-carbon sources *S. cerevisiae* DGI342 batch simulations using Yeast8 (A-C), ecYeast8 (D-F) and ecYeast8 with additional regulation (G-H) compared to experimental data (D,) [27]. Colored areas represent different phases in the process: glucose consumption (orange), sucrose hydrolysis and glucose consumption (green), fructose consumption (red) and mannose consumption (purple).

**Figure B.8:**
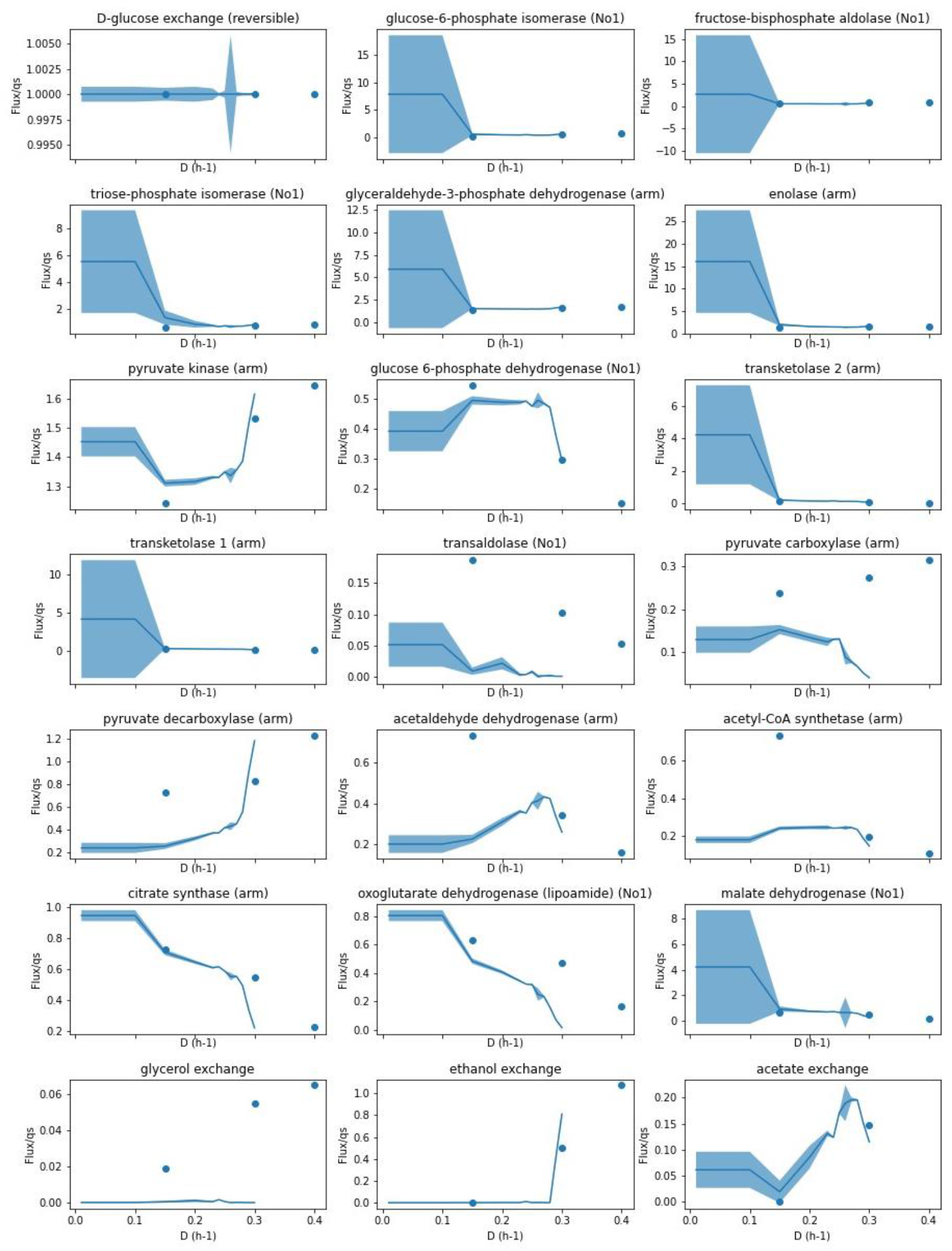
Comparison of intracellular flux predicted by ecYeast8 using flux sampling (-) to fluxes measured with ^13^*C* flux analysis ( ) at different growth rates (D) [30]. The median ± MAD of 10000 samples is shown.

